# CSF1R inhibition by small molecule affects T-helper cell differentiation independently of microglia depletion

**DOI:** 10.1101/2021.12.21.473532

**Authors:** Fengyang Lei, Naiwen Cui, Chengxin Zhou, James Chodosh, Demetrios G. Vavvas, Eleftherios I. Paschalis

## Abstract

Colony-stimulating factor 1 receptor (CSF1R) inhibition has been proposed as a specific method for microglia depletion. However, recent work revealed that in addition to microglia, CSF1R inhibition also affects other innate immune cells, such as peripheral monocytes and tissue-resident macrophages of the lung, liver, spleen, and peritoneum. Here, we show that this effect is not restricted to innate immune cells only, but extends to the adaptive immune compartment. CSF1R inhibition alters the transcriptional profile of bone marrow cells that control T helper cell activation. *In vivo* or *ex vivo* inhibition of CSF1R profoundly changes the transcriptional profile of CD4^+^ cells and suppresses Th1 and Th2 differentiation in directionally stimulated and unstimulated cells and independently of microglia depletion. Given that T cells also contribute in CNS pathology, these effects may have practical implications in the interpretation of relevant experimental data.

**Significance statement:** Here we show that CSF1R inhibition affects not only innate immune cells and microglia, but also the adaptive immune compartment by suppressing Th1/Th2 differentiation of CD4^+^ cells, independently of microglia depletion.

## Introduction

We recently showed that colony stimulating factor 1 receptor (CSF1R) inhibition not only affects microglia but also alters the function of bone marrow-derived macrophages(1, 2); confirmed by others using CSF1R knock-out mice(3, 4). However, it is still unknown if these off-target effects extend to other cells that belong to the adaptive immune compartment. This is critical since recent studies have demonstrated the role of T cells in CNS pathology, especially in models of autoimmune diseases and glaucoma(5–7).

Here, we adopt *in vivo* and *ex vivo* models, along with targeted transcriptomics and Ingenuity Pathway Analysis (IPA), to show that CSF1R inhibition not only affects innate immune cells, but also pathways in the bone marrow that control T-cell activation. In particular, CSF1R inhibition causes overexpression of *Hdac9*, *Cd1d*, *Lrmp*, *Pax5*, *Pecam1*, *Cd8A*, *Stim2*, *Blnk*, *B2m*, *Rbpj* and repression of *Spp1* genes in bone marrow cells **(fig. 1 A, B)**. Cessation of the inhibitor for 1 month does not restore these changes, *Nocoa6*, *Cd8a*, *Cd14*, *Cd3g*, *Stim2*, *B2m*, *Pecam1*, *Hdac9*, *Spp1* become over expressed and *Hdac4*, *CSF2*, *Nos2*, *Tal1 and Notch4* repressed **(fig. 1 A, B)**. *Spp1* is the single gene that switches from repression to overexpression after cessation of the inhibition **(fig. 1 B)**, while *interleukin 10* (*Il-10*), *nuclear receptor coactivator 6* (*Ncoa6*) and *CD14* **(fig. 1 B)** are the most differentially expressed genes immediately, 1 and 2 months after cessation of the inhibitor. Principal Component Analysis (PCA) of the gene array confirms that CSF1R inhibition alters the transcriptional profile of hematopoiesis for the long-term **(fig. 1 D)**. Further analysis using ingenuity pathway predicts that CSF1R inhibition leads to upregulation of colony stimulating factor 2 (CSF2), which alters canonical pathways associated with Th1 and Th2 activation **(fig. 1 E)**.

**Figure 1.**
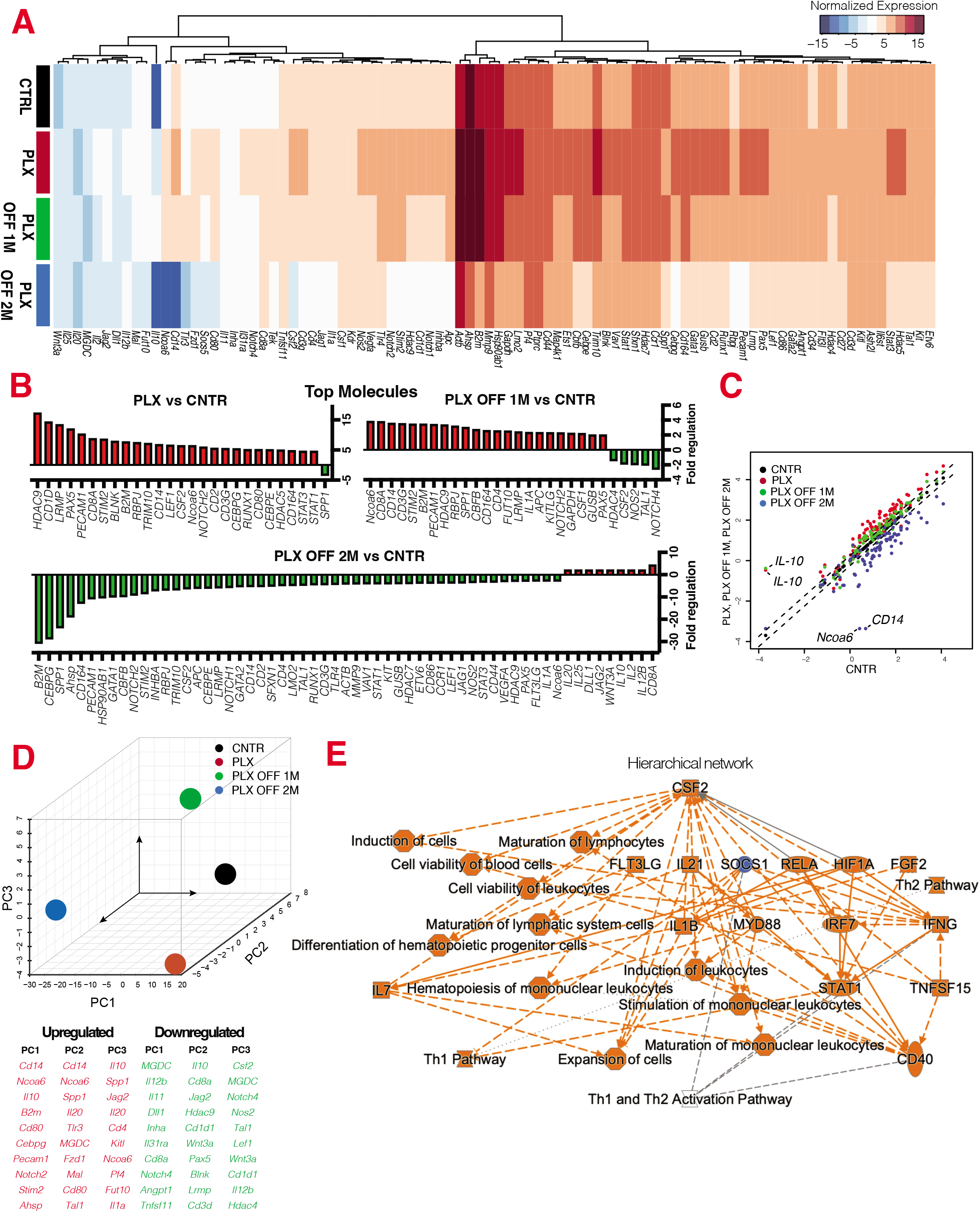
CSF1R inhibition alters the transcriptional profile of bone marrow cells and that control Th1/Th2 activation. Gene expression analysis of bone marrow cells isolated from C57BL/6J mice using targeted transcriptomics for common hematopoietic genes immediately, 1 and 2 months after cessation of CSF1R inhibition. (**A, B**) CSF1R inhibition causes a wide range of transcriptional changes in hematopoietic gene expression, (**C**) especially through the deregulation of *Il-10*, *Cd14* and *Ncoa6*. (**D**) Principal component analysis suggests that CSF1R inhibition causes long-term transcriptional changes in bone marrow cells, even 2 months after cessation of the inhibitor. (**E**) Ingenuity pathway analysis of the gene array predicts that CSF1R inhibition upregulates CSF2, which alters the regulation of canonical pathways that control Th1 and Th2 activation. **(A-D)** n=5 per group. All entities have p-value ≤ 0.05, regulators and processes have |z-score| ≥ 2, and regulators are limited to genes only.

The predicted model from IPA was confirmed *in vivo*, by acquiring CD4^+^ cells from PLX5622 treated (fed) mice for targeted transcriptomics of T helper cell differentiation. CSF1R inhibition for 3 weeks profoundly affects T helper cell differentiation transcriptional profile, notably through suppression of interleukin 12 receptor subunit beta 2 (Il12rb2) and over expression of CSF2 **(fig. 2 A)**. Further analysis with ingenuity pathway predicts that these gene changes affect Th1 and Th2 cell activation **(fig. 2 B)**. Indeed, inhibition of CSF1R causes dosedependent reduction in CD4^+^ cell survival and suppression of Th1/Th2 differentiation *ex vivo* **(fig. 2 C),** independently of microglia depletion. Cessation of the inhibitor restores Th1/Th2 differentiation, IL-12 or IL-4 stimulation enhances Th1/Th2 differentiation, respectively, in PLX5622 fed mice **(fig. 2 D-K)**, though mice fed with PLX5622 have lower number of CD4^+^ cells **(fig. 2 E)**.

**Figure 2.**
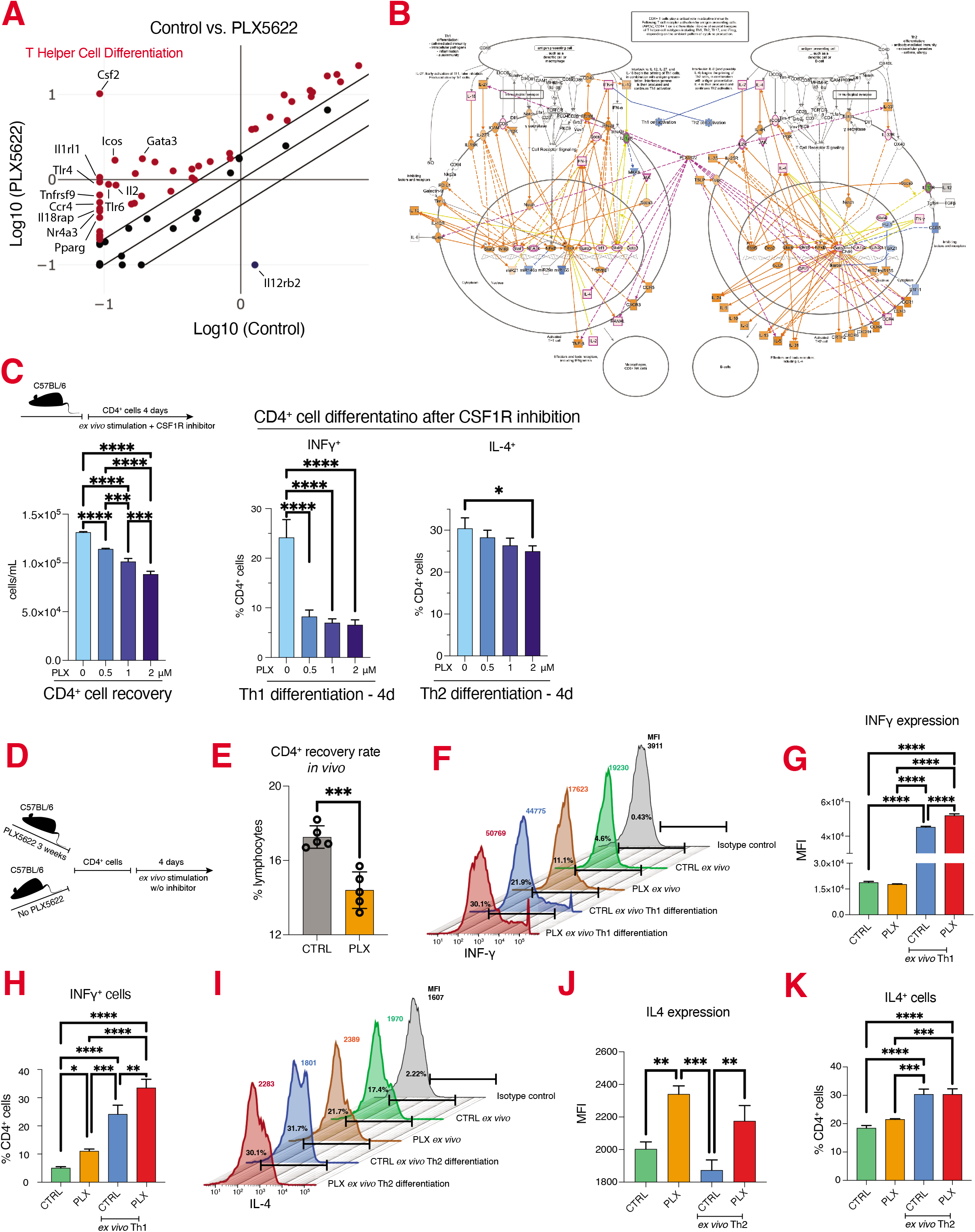
CSF1R inhibition alters the transcriptional profile CD4^+^ cells and suppressed Th1 and Th2 differentiation. **(A)** Targeted transcriptomics on CD4^+^ cells reveals that CSF1R inhibition causes significant changes in genes that control T helper cell differentiation. **(B)** Ingenuity pathway analysis predicts that these gene changes suppress Th1 and Th2 activation pathways. **(C)** CSF1R inhibition ex vivo causes dose dependent reduction in CD4^+^ cell number and suppression of Th1/Th2 differentiation. (**D**) Schematic representation of the experimental setup to assess Th1/Th2 differentiation after cessation of CSF1R inhibitor. **(E)** Percentage of CD4^+^ cells recovered from control and PLX treated mice. Histogram, mean fluorescent intensity and percentage of CD4^+^ cells that express **(F-H)** INFγ (Th1) or **(I-K)** Il-4 (Th2). **(F-K)** Although mice treated with CSF1R inhibition have lower numbers of CD4^+^ cells, cessation of the inhibitor *ex vivo* restores their ability to differentiate, though directional Th1/Th2 stimulation ex vivo enhances the response.

Previous studies have assumed that CSF1R inhibition affects only microglia(8–11). However, CSF1R inhibition was also shown to affect peripheral monocytes and the function of macrophages(1–4). Here, we provide insights into the role of CSF1R inhibition on the adaptive immune compartment and show that it causes changes that affect CD4^+^ cells and T helper cell differentiation. Considering that T cells have been recently shown to participate in CNS pathologies, especially in autoimmune conditions and glaucoma, future experiment data with CSF1R inhibitors need to account for these changes in the interpretation of relevant experimental data.

## Materials and Methods

### Mouse model

Animal experiments were performed in accordance with the Association for Research in Vision and Ophthalmology Statement for the Use of Animals in Ophthalmic and Vision Research, and the National Institutes of Health (NIH) Guidance for the Care and Use of Laboratory Animals. This study was approved by the Mass. Eye and Ear Animal Care Committee. Mice at 6-8 weeks old of both genders were used: C57BL/6J (Jackson Laboratory, Stock#: 000664). PLX5622 (PLexxikon, Inc) was formulated into AIN-76A chow (Research Diets, Inc) at the dose of 1200 ppm and given *ad libitum* for 3 weeks.

### *Ex vivo* differentiation

Spleen and lymph nodes from C57BL/6 mice were surgically removed and single cells were obtained mechanically using a 70μm cell strainer (BD Falcon). Red blood cells were lysed by ACK lysis buffer (Lonza) before CD4^+^ isolation, achieved using the CD4+ T cell isolation Kit (Miltenyi) and according to manufacturer’s instructions. 0.5 ×10^6^ CD4^+^ cells/mL were suspended in 10% FBS RPMI-1640 (ThermoFisher) supplemented with 5 ng/mL IL-2 (PeproTech) and 0.5 μg/mL anti-CD28 antibody (clone:37.51, BioLegend) and seeded into 2 μg/mL anti-CD3 antibody (clone: 145-2C11, BioLegend) pre-coated 24-well plates. Th1 differentiation was achieved using 1 μg/mL anti-IL-4 antibody (clone: 11B11, Biolegend), 5 ng/mL IL-2 (PeproTech), and 10 ng/mL IL-12 (PeproTech) in the culture media. Th2 differentiation was achieved using 1 μg/mL anti-IFN-γ antibodies (clone: XMG1.2, Biolegend), 5 ng/mL IL-2(PeproTech), and 10 ng/mL IL-4 (PeproTech) in the culture media. Cells were incubated for 96 hours in 5% CO2 at 37°C. Cell Activation Cocktail with Brefeldin A (Biolegend) was added to the culture media 5 hours prior to harvesting and staining the cells. For PLX5622 treatment, different concentrations of PLX5622 were added to the culture media during the stage of differentiation to Th1/Th2.

### Flow cytometry

Cultured CD4^+^ T cells and *ex vivo* splenocytes were processed for flow cytometry. Anti-murine CD16/32 (Clone: 2.4G2, eBiosciences) was used for blocking. Anti-CD3 (Clone: 17A2), anti-CD4 (Clone: GK1.5), anti-IL-4 (Clone: 11B11), anti-INF-γ (Clone: XMG1.2) were purchased from BioLegend. Data were acquired by LSRII cytometer (BD Biosciences) and analyzed by FLowJo V10 (Tree Star).

### RT2 profiler assay

A total of 1 μg of RNA was used for each array analysis. cDNA was synthesized by RT2 First Strand Kit (Qiagen, Hilden, Germany) and then mixed with RT2 SYBR Green ROX qPCR Mastermix (Qiagen) before loading into the array well (Mouse hematopoiesis: PAMM-054ZA; Mouse T helper cell differentiation: PAMM-503ZA and Mouser Retinoic Acid Signaling: PAMM-180ZA). qPCR was performed by using QuantStudio 3 (ThermoFisher) according to manufacturer’s manual. Data analysis was performed by using Qiagen’s data analysis web portal (https://geneglobe.qiagen.com/us/analyze/) to generate a normalized expression profile. Further normalization was performed based on 2^ (-Delta Delta CT). Gene expression heat-map, 3D PCA plot, and scatter plot were generated using the normalized gene set and built-in functions of R-software, per previous instructions (Maechler, M., Rousseeuw, P., Struyf, A., Hubert, M., Hornik, K. (2013). Cluster: Cluster Analysis Basics and Extensions. R package version 1.14.4.).

### IPA analysis

Further analysis of the gene set was performed using Ingenuity Pathway Analysis (IPA, Redwood City, CA, USA) that assesses biological networks and disease functions associated with the gene profiles. Comparative Ingenuity Pathway Analysis (IPA) (http://www.ingenuity.com) was also performed to assess differentially regulated pathways, upstream regulators, and disease/functions associated to CSF1R inhibition. IPA uses a manually curated database which contains information from the published literature as well as many other sources, including several gene expression and gene annotation databases such as IntACT, BIND, MiPs et al.(12) A threshold of 2-fold change in gene expression and P ≤ 0.05 was set for every analysis. Data were ranked based on their z-score.

### Statistical analysis

Data were analyzed with GraphPad (Prism 2.8.1, San Diego, CA) using two-tailed unpaired t-test and ordinary one-way ANOVA with Dunnet’s correction for multiple comparisons. Statistical significance was set at P ≤ 0.05.

## Author contributions

FL designed experiments, acquired data and analyzed data; NC, CZ analyzed data; DGV wrote and reviewed the manuscript; JC reviewed the manuscript; EIP designed experiments, analyzed data, and wrote the manuscript.

## Data Availability

All data necessary for study replication have been included in the submission. Materials are available commercially and are listed in Materials and Methods.

## Acknowledgments

This work was supported by the Boston Keratoprosthesis Research Fund, DoD 2019A016948, and core grant P30EY003790; PLX5622 was kindly provided by Plexxikon Inc. We would like to thank Dr. Thomas Dohlman for providing columns for T-cell isolation.

